# Mis-expression of the Alzheimer’s disease associated gene Ankyrin causes memory loss and shortened lifespan in *Drosophila*

**DOI:** 10.1101/423129

**Authors:** James P Higham, Bilal R Malik, Edgar Buhl, Jenny Dawson, Anna S Ogier, Katie Lunnon, James JL Hodge

**Author notes:** These authors contributed equally to this study (James P Higham, Bilal R Malik). corresponding author: Dr. James Hodge.

## Abstract

Alzheimer’s disease (AD) is the most common form of dementia and is characterized by the accumulation of extracellular amyloid beta (Aβ) plaques and intracellular neurofibrillary tangles of hyperphosphorylated Tau, including the 4R0N isoform. Recent epigenome-wide association studies (EWAS) of AD have identified a number of loci that are differentially methylated in AD cortex. Indeed, hypermethylation of the *Ankyrin 1* (*ANK1*) gene in AD has been reported in the cortex in numerous different post-mortem brain cohorts. Little is known about the normal function of ANK1 in the healthy brain, nor the role it may play in AD. We have generated *Drosophila* models to allow us to functionally characterize *Drosophila Ank2*, the ortholog of human *ANK1*. These models have targeted reduction in the expression of *Ank2* in neurons. We find that *Drosophila* with reduced neuronal *Ank2* expression have shortened lifespan, reduced locomotion, reduced memory and reduced neuronal excitability similar to flies overexpressing either human mutant *APP* (that leads to Aβ42 production) and *MAPT* (that leads to 0N4R Tau). Therefore, we show that the mis-expression of *Ank2* can drive disease relevant processes and phenocopy some features of AD and we propose targeting ANK1 may have therapeutic potential. This represents the first study to characterize a gene implicated in AD, which was nominated from EWAS.

**Author summary:** The majority (>95%) of Alzheimer’s disease (AD) cases are sporadic, with their incidence attributed to common genetic mutations, epigenetic variation, aging and the environment. There is no cure for AD and only limited treatment options which only treat the symptoms of AD and only work in some people. Recent epigenome-wide association studies (EWAS) in AD have highlighted hypermethylation of the Ankyrin1 (*ANK1*) gene in AD cortex. Little is known of the normal role of the gene in the brain. Here, we have demonstrated that *Drosophila* with reduced neuronal expression of the *Drosophila* ortholog of human *ANK1 (Ank2)*, can drive AD relevant processes including locomotor difficulties, memory loss and shortened lifespan similar to expression of human amyloid-Beta or tau mutant proteins. Furthermore, increasing *Ank2* expression reversed the memory loss caused by expression of human amyloid-Beta or tau mutant proteins, suggesting that targeting *ANK1* may have therapeutic potential. This represents the first study to characterize a gene implicated in AD, which was nominated from EWAS.

## INTRODUCTION

AD is the most common form of dementia, with patients suffering from premature death and accelerated cognitive decline including memory loss. Post-mortem examination of AD brain samples reveals the accumulation of extracellular amyloid-beta (Aβ) plaques and intracellular neurofibrillary tangles (NFTs) of hyperphosphorylated microtubule associated protein Tau (MAPT), which is accompanied by gliosis, neuronal cell loss and brain atrophy. Aβ is produced by the amyloidogenic cleavage of the amyloid precursor protein (APP) gene by β and γ secretases resulting in the formation of neurotoxic aggregating Aβ peptides with the 42 amino acid (aa) peptide (Aβ42) being more toxic than Aβ40 [1]. Tau exists in six different isoforms, which vary with the number of C-terminal aggregating tubulin binding repeats (3R or 4R) and the number of N-terminal domains (0N, 1N or 2N) [2]. The 4R isoforms are upregulated in AD brain and show stronger tubulin binding and aggregation than the 3R isoforms [2]. Another contributing factor to Tau aggregation is its hyperphosphorylation by a number of different kinases [2, 3]. Less than 5% of AD cases are due to autosomal dominant mutations in the amyloid precursor protein (*APP*), presenilin 1 (*PSEN1*) or *PSEN2* genes. The remainder of AD cases are sporadic, with incidence attributed to both genetic and environmental risk factors. Genome-wide association studies (GWAS) have identified a number of genes where common genetic variation is associated with increased risk of sporadic AD, including *APOE* (ε4 allele), *BIN1* and *PICALM* genes amongst others [4] with many additional risk factors for AD being associated with lifestyle and/or the environment.

Epigenetics refers to the mitotically and meiotically heritable changes in gene expression without alterations in the underlying DNA sequence for instance by DNA methylation and downregulation of genes [5]. This also potentially allows for alterations in gene expression in response to environmental variation, such as stress, diet or exposure to environmental chemicals. In order, to characterize the contribution of epigenetic mechanisms to AD etiology, recent epigenome-wide association studies (EWAS) of AD [6-8] have been performed and have identified a number of genetic loci that are associated with increased risk of AD. One locus that showed consistent cortical AD-associated hypermethylation in five independent cohorts resided in the *ANK1* gene [6, 7, 9]. A recent publication has demonstrated that *ANK1* DNA methylation in entorhinal cortex is observed in only certain neurodegenerative diseases [10]. This study reported disease-associated hypermethylation in AD, Huntington’s disease (HD) and to a lesser extent Parkinson’s disease (PD). The authors showed that disease-associated hypermethylation was only seen in donors with vascular dementia (VaD) and dementia with Lewy bodies (DLB) when individuals had co-existing AD pathology; in individuals with “pure” VaD or DLB, no *ANK1* hypermethylation was observed. ANK1 is an integral membrane and adaptor protein, that mediates the attachment of membrane proteins such as ion channels, cell adhesion proteins and receptors with the spectrin-actin cytoskeleton and is important for cell proliferation, mobility, activation, and maintenance of specialized membrane domains [11].

Most of our understanding of the molecular changes that cause AD pathology comes largely from experiments using rodents to model genetic variation; however, these are models of familial AD, and do not recapitulate sporadic disease. As such, new drugs effective in these rodents have not translated to any new successful treatments for AD and highlighting the need to generate and characterize new models of sporadic AD [6, 12-16]. For example, widespread constitutive overexpression of a range of different mammalian *APP* transgenes in rodents lead to a range of AD relevant phenotypes such as shortened lifespan, movement and memory deficits however by itself leads to little or no neurodegeneration or development of NFTs [13, 17, 18]. Likewise, widespread constitutive overexpression of a range of different mammalian *MAPT* transgenes in rodents lead to a range of dementia relevant phenotypes, but little neurodegeneration [3, 19, 20]. However, much of this work characterized rodents overexpressing P301L mutant *MAPT*, which is associated with frontotemporal dementia (FTD) and not AD [21, 22] or with mice expressing multiple mutant transgenes from different diseases [23]. It is not currently known how Aβ and Tau pathology are linked, however the amyloid cascade hypothesis suggests that Aβ pathology leads to the other hallmarks of AD, including the spread of NFTs [1] and recent work suggests changes in neuronal excitability and calcium (Ca^2+^) signaling may be important for their connection and disease progression [24, 25], but exactly how remains unknown.

In *Drosophila*, neuronal overexpression of different human APP products (including Aβ42) and mutants has been reported to cause degeneration of the photoreceptor neurons of the fly eye, shortened lifespan, change in neuronal excitability as well as movement, circadian, sleep and learning defects in a number of different studies [26-32]. Likewise, neuronal overexpression of human Tau isoforms associated with AD have been shown to result in degeneration of the photoreceptor neurons of the fly eye, shortened lifespan, movement and learning defects in many different studies [33-41]. Fewer animal models have determined the effect of expression of human Aβ and Tau together [39, 40, 42, 43], a scenario that more accurately reflects the progression of AD in humans [1, 2, 24, 44-46]. Furthermore, fewer studies have compared the effect of common variants nominated from GWAS for AD [43, 47-50] and to our knowledge none from EWAS for AD. Thus, many genomic and epigenomic loci nominated in these studies remain uncharacterized in any living organism, with many of these genes or loci having a completely unknown function in the brain [6, 46, 49-52].

In order to address these issues, we have generated and characterized the first animal model to investigate the function of a locus nominated from EWAS in AD. We have investigated *ANK* mis-expression and compared its AD relevant phenotypes to *Drosophila* models expressing either (a) human mutant *APP* (which results in an aggregating form of oligomerized Aβ42), (b) MAPT (resulting in 0N4R Tau), (c) *APP* (Aβ42) with *MAPT* (0N4R Tau) and (d) *APP* (Aβ42) or *MAPT* (0N4R Tau) with *Ank* mis-expression, finding that mis-expression of these AD associated genes cause similar reduction in lifespan, movement, memory and neuronal excitability.

## RESULTS

### Mis-expression of human mutant APP, MAPT or Ank2 shortens lifespan

Increased neuronal levels of Aβ or Tau lead to AD pathology and early death [1, 45]. We therefore overexpressed in all neurons either an aggregating form of human mutant *APP* that encodes oligomerized human Aβ42 (tandem Aβ42 [29]) or human MAPT (0N4R Tau [35, 37]) both of which caused premature death reducing the flies’ lifespan by about 25% (Fig. 1). In order to test for additive, synergistic or no change in effect we overexpressed both mutant *APP* (Aβ42) and *MAPT* (0N4R) together in all neurons and got a further reduction to exactly half the lifespan of a normal fly. The reduction in median lifespan due to co-expression was equivalent to an additive effect of the shortening of life due to mutant *APP* (Aβ42) and *MAPT* (0N4R) alone, this suggests that Aβ42 and Tau pathology may act in separate pathways to cause neurotoxicity and early death.

**Figure 1.**
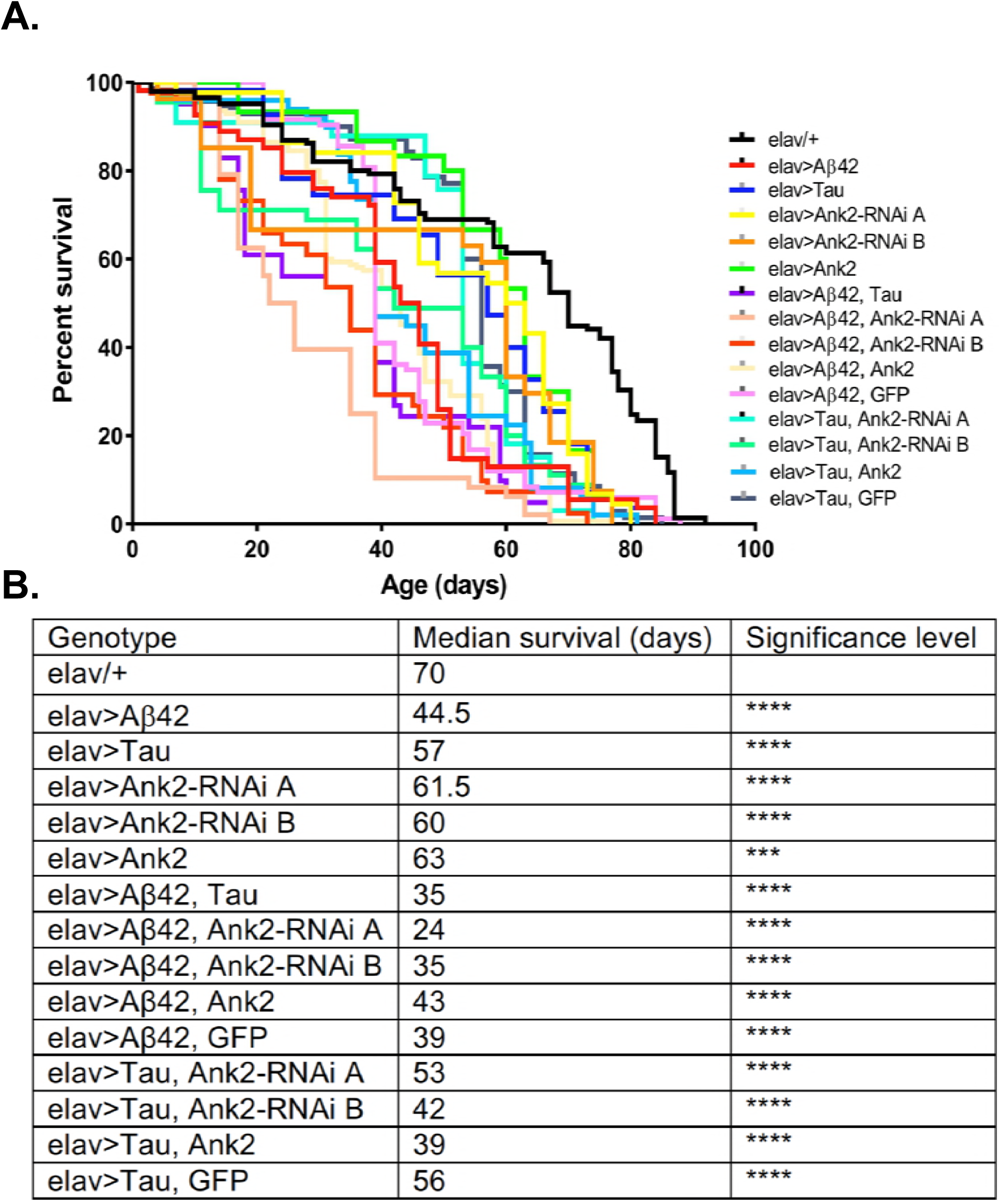
The effect of expression of human mutant *APP* (Aβ42), *MAPT* (Tau 0N4R) and *Drosophila Ank2* on *Drosophila* lifespan. **(A)** Survival curves of pan-neuronal expressing Aβ42 (*elav>Aβ42*), Tau (*elav>Tau*), Ank2-RNAi (*elav>Ank2-RNAi line A or B*), Ank2 (*elav>Ank2*), Tau with Aβ42 (*elav>Tau, Aβ42*), Aβ42 with *Ank2-RNAi* (*elav>Aβ42, Ank2-RNAi line A or B*), Aβ42 with Ank2 (*elav>Aβ42, Ank2*), Aβ42 with Ank2 (*elav>Aβ42, Ank2*), Aβ42 with Ank2 (*elav>Aβ42, GFP*), Tau with *Ank2-RNAi* (*elav>Tau, Ank2-RNAi line A or B*), Tau with Ank2 (*elav>Tau, Ank2*), Tau with GFP (*elav>Aβ42, GFP*) compared to wild type control (*elav/+*) flies kept at 25°C. Misexpression of all AD genes caused a significant reduction in lifespan compared to control using the Kaplan-Meier and log rank test (n>100 per genotype of flies). **(B)** Table listing all genotypes characterized with median lifespan (days) and significant reductions in lifespan as determined by Kaplan-Meier and log rank test and indicated as *p<0.05, **p<0.01, ***p<0.001, ****p<0.0001 and used in all subsequent figures.

Recently, human *ANK1* gene has been shown to be differentially methylated in AD, with some evidence for altered expression of some transcript variants in AD [7]. There are two other *ANK* genes in the human genome, *ANK2* and *ANK3*. In *Drosophila,* there are two *Ank* genes, the ubiquitously expressed *Ank1*, while *Ank2* is specifically expressed in neurons [53-57]. The closest orthologue to *Drosophila Ank1* is human *ANK3* with 51% total amino acid identity while the closest homolog of *Drosophila Ank2* is human *ANK1* with 43% amino acid identity. Human *ANK2* is more similar to *Drosophila Ank1* (49% identity) than *Drosophila Ank2* (33% identity). Because hypermethylation of human *ANK1* has been reported in AD cortex [7], we used *RNAi* to knock-down expression of *Ank1* and *Ank2* comparing the effect of reducing these genes in all *Drosophila* neurons using two different *RNAi* transgenes designed to non-overlapping regions of each gene. Pan-neural mis-expression of *Drosophila Ank1* did not cause any AD relevant behavioral deficits (Fig. 2A-B), while reduction in *Ank2* using the same promoter caused both a reduction in locomotion (Fig. 4) and 1 hour memory (Fig. 2B). Therefore, we conclude, that *Drosophila Ank2* is the closest functional ortholog of human *ANK1*, which we characterized in subsequent experiments.

**Figure 2.**
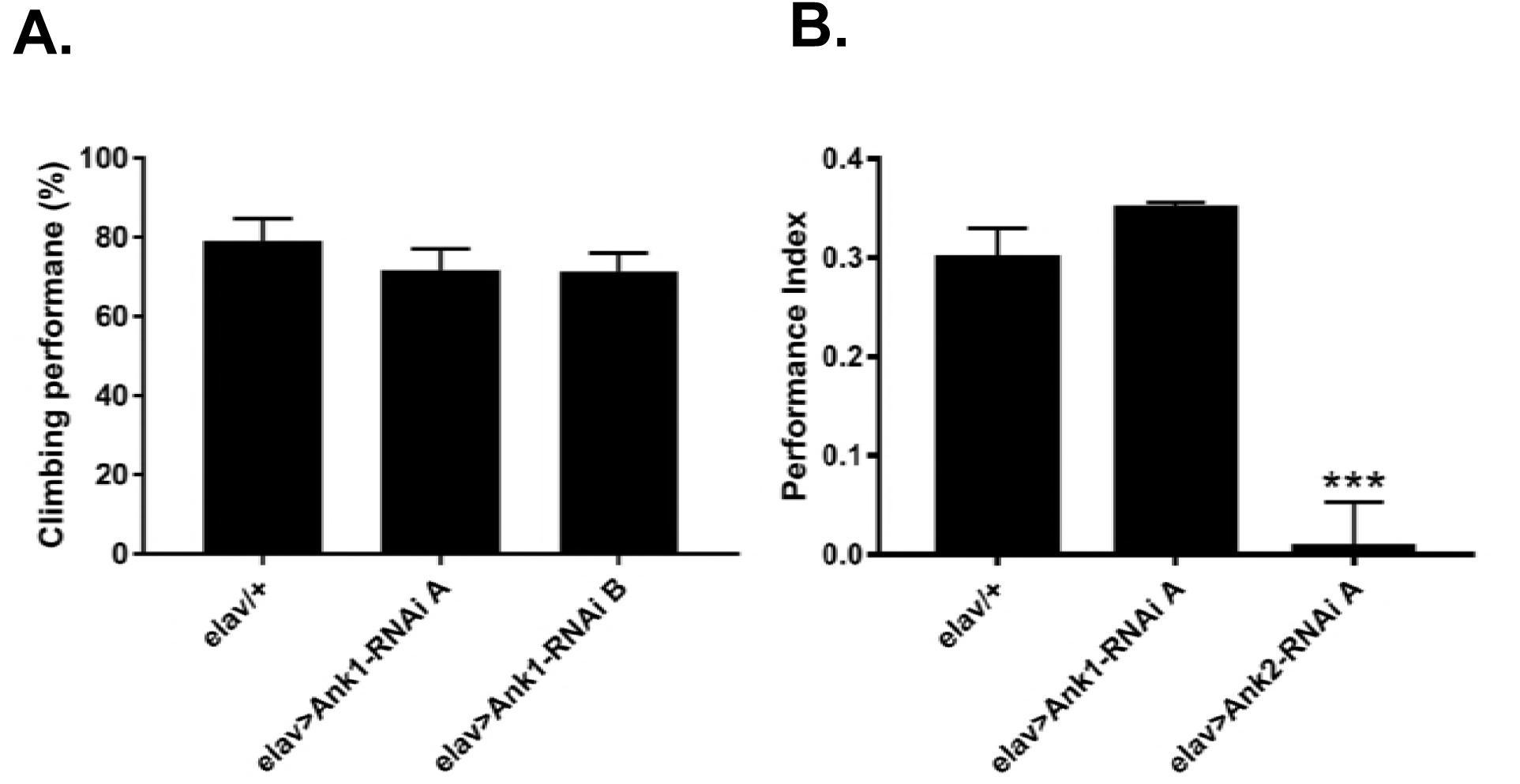
The effect of pan-neuronal expression of *Drosophila Ank1* and *Ank2* on behavior (**A**) The negative geotaxis-climbing reflex was used to quantify motor deficits. Pan-neuronal reduction of *Ank1* (*elav-Gal4>Ank1-RNAi line A or B*) was compared to control using one-way ANOVA with Dunnett’s multiple comparison and showed no significant difference. n≥6 groups of ten flies. (**B**) 1 hour memory was assessed using the olfactory shock-conditioning assay. Pan-neuronal reduction in Ank2 (*elav>Ank2-RNAi line A*) as opposed to Ank1 (*elav>Ank1-RNAi line A*) caused a significant reduction in memory compared to control using one-way ANOVA with Dunnett’s multiple comparisons. Error bars are standard error of the mean (SEM). n≥4 groups of ∼50 flies.

We found that pan-neural reduction of *Ank2* resulted in a shortening of lifespan by 15% compared to control (Fig.1). We verified are results by using two independent *RNAi* lines (A and B) to non-overlapping sequences in *Ank2*, both lines gave similar results in all assays. Depending on the genomic location of DNA methylation, it can result in either increased or decreased expression of the target gene [58]; we therefore tested if neuronal *Ank2* overexpression affected longevity and found that it also shortened lifespan (∼10%), but to a slightly less extent. In order to test if there was a genetic interaction between mutant *APP* (Aβ42) or *MAPT* (0N4R) and *Ank2* we generated flies that co-expressed mutant *APP* (Aβ42) or *MAPT* (0N4R) with the *Ank2* transgenes. We found that the detrimental effect of each gene on lifespan was additive, again suggesting the molecules may act in separate senescence pathways. One potential cause of a reduction or suppression of toxicity of a gene product is due to a dilution of Gal4. Whereby, a single Gal4 transcription factor driving expression by binding to a single UAS transgene (e.g. mutant *APP* (Aβ42) or *MAPT* (0N4R)) gene gets diluted when adding a second UAS site of a gene (e.g. *Ank2*) is present. This results in a reduction in the amount of the AD toxic gene product being made giving a false positive of a suppression phenotype. To control for such a dilution effect, we generated flies that co-expressed a second unrelated neutral gene product (GFP) with either human mutant *APP* (Aβ42) or *MAPT* (0N4R Tau). We found no reduction in the toxicity of the transgenes in terms of shortening lifespan and in any other AD relevant phenotypes tested (i.e. neurodegeneration, locomotion or memory), showing any suppression of neurotoxicity caused by human mutant *APP* (Aβ42) or *MAPT* (0N4R Tau) overexpression was likely due to *Ank2* overexpression.

### Overexpression of human mutant APP or mutant MAPT but not mis-expression of Ank2 caused degeneration of photoreceptor neurons

Increased levels of aggregating toxic Aβ and Tau lead to neurodegeneration and AD. In order to model this neurotoxicity in *Drosophila*, human mutant *APP* causing Aβ42 production (Fig. 3B) or mutant MAPT (0N4R Tau) (Fig. 3C) was expressed throughout development and adulthood in the photoreceptor neurons of the eye, resulting in a so called “rough eye” phenotype, where the degeneration and loss of the normally regularly arrayed ommatidia of the compound eye of wild-type flies (Fig. 3A) gives rise to a disorganized and smaller eye. The loss of photoreceptors could be quantified by measuring the total surface area of the eye, which showed that expression of human mutant *APP* (Aβ42) and *MAPT* (0N4R Tau) reduced the size of the eye by 37% and 40%, respectively (Fig. 3H). This degenerative phenotype became only slightly worse when both human mutant *APP* (Aβ42) and *MAPT* (0N4R Tau) were co-expressed (Fig. 3D), resulting in a 53% reduction in eye size, suggesting that Aβ42 and Tau act in partially overlapping pathways to cause neurotoxicity in the fly eye. Reduction (Fig. 3E-F) or overexpression (Fig. 3G) of *Ank2* alone did not cause degeneration or reduction in size of the eye (Fig. 3H). Overexpression of *Ank2* with human mutant *MAPT* (0N4R Tau) rescued the reduction in size of eye caused by human mutant *MAPT* (0N4R Tau) alone to levels similar to wild-type, while co-expression of human mutant *MAPT* (0N4R Tau) with reduced *Ank2* continued to result in a significantly reduced eye size.

**Figure 3.**
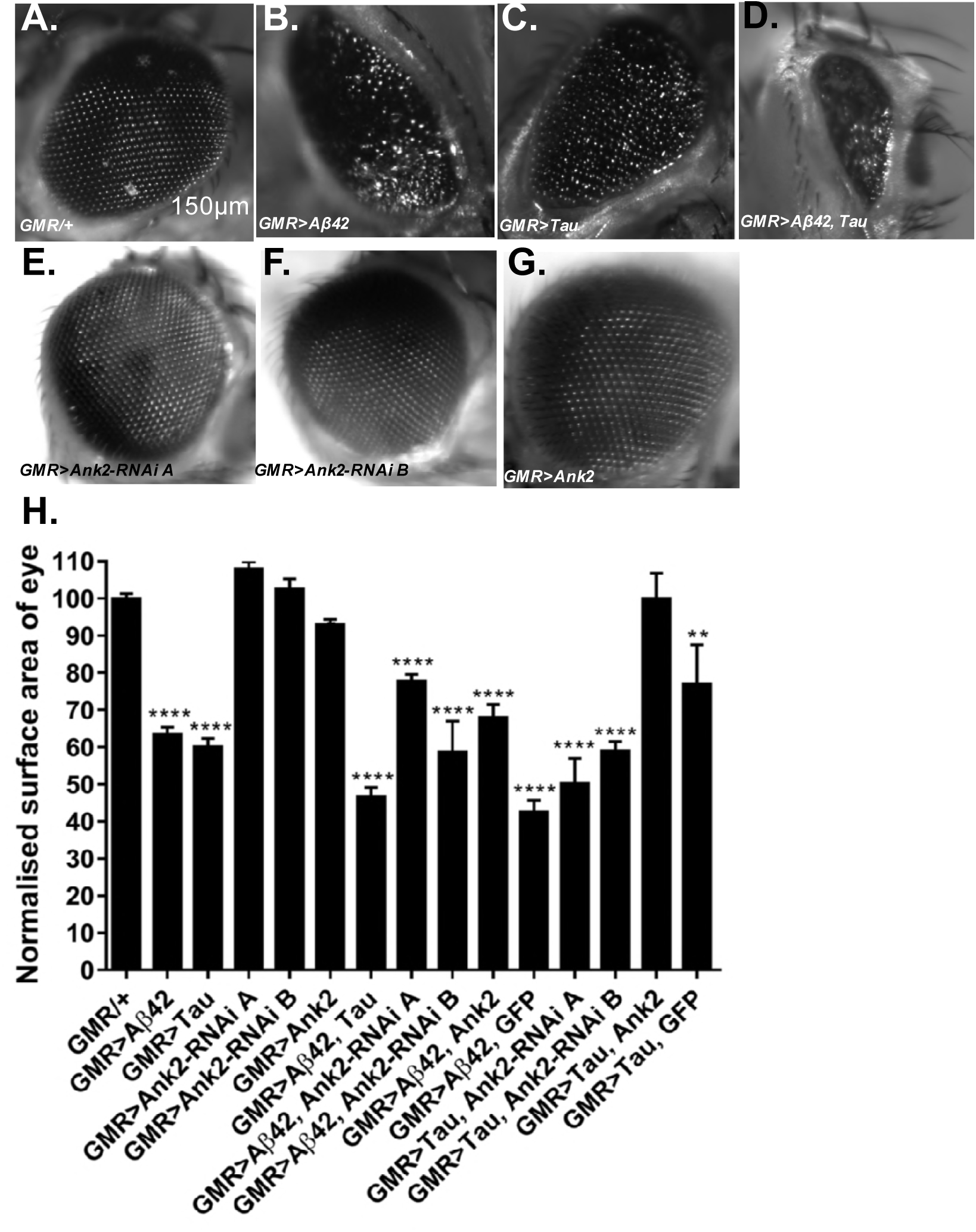
The effect of expression of human mutant *APP* (Aβ42), *MAPT* (Tau 0N4R) and *Drosophila Ank2* on degeneration of photoreceptor neurons. Images of compound eyes of (**A**) control fly (*GMR-Gal4/+*) showing the regular alignment of ommatidia compared to photoreceptors neurons overexpressing (**B**) Aβ42 (*GMR-Gal4>Aβ42*), (**C**) Tau (*GMR-Gal4>Tau*) and (**D**) Tau with Aβ42 (*GMR-Gal4>Tau, Aβ42*) which were smaller and displayed a “rough eye” phenotype. (**D-E**) photoreceptors expressing *Ank2-RNAi* (*GMR>Ank2-RNAi line A or B*) or (**F**) Ank2 (*GMR>Ank2*) appeared normal. (**H**) Degeneration of photoreceptor neurons was quantified as normalized percentage surface area of the eye of genotypes compared to the mean of the control (*GMR/+*) which was set at 100%, comparisons were made between to mutant genotypes using one-way ANOVA with Dunnett’s multiple comparisons. Genotypes that included expression of *APP* (Aβ42) or *MAPT* (Tau 0N4R) in the eye showed a significantly reduced size of eye, while those mis-expressing and *Drosophila Ank2* did not. Co-expression of *MAPT* (Tau 0N4R) and *Ank2* rescued eye size to a level indistinguishable from wildtype. Error bars are SEM. n≥7 eyes per genotype.

**Figure 4.**
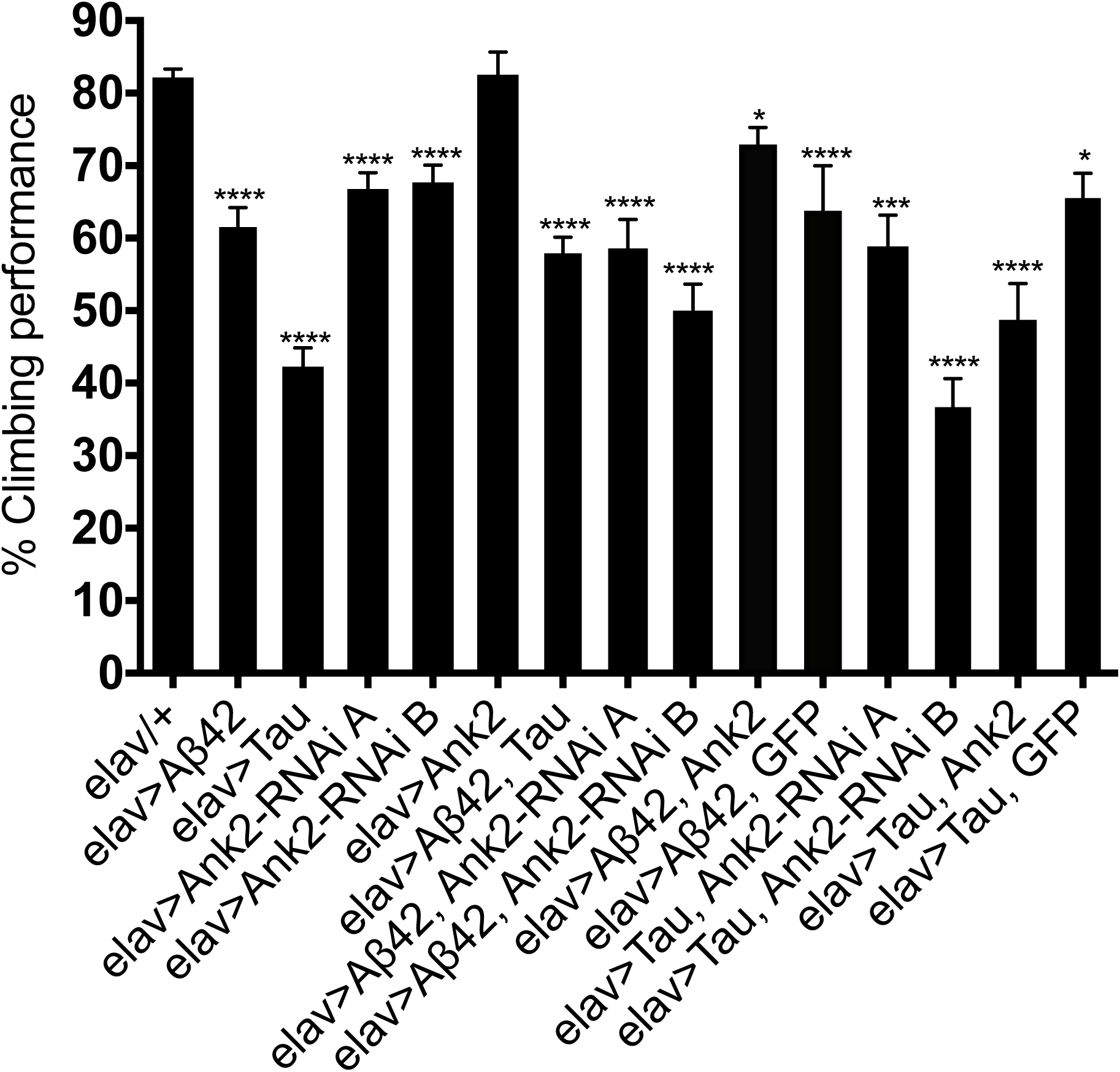
The effect of expression of human mutant *APP* (Aβ42), *MAPT* (Tau 0N4R) or *Drosophila Ank2* on locomotor behavior. The negative geotaxis-climbing reflex was used to quantify locomotor behavior. About 80% of control (*elav-Gal4/+*) flies were able to climb to the top of test vial within 10 seconds this compared to experimental genotypes expressing AD transgenes throughout their nervous system. Mis-expression of all AD associated genes except for overexpression of *Ank2* alone, resulted a significant reduction in climbing performance using one-way ANOVA with Dunnett’s multiple comparisons. Error bars are SEM. n≥15 groups of ten flies.

### Overexpression of human mutant APP and MAPT genes and reduction in Ank2 expression caused locomotor deficits

Flies show a negative geotaxis reflex, such that after tapping on a surface about 80% of young control flies will climb to the top of a tube in 10 seconds, indicative of a healthy coordinated nervous system (Fig. 4). In order to quantify any detrimental effect of *human mutant APP* (Aβ42) and *MAPT* (0N4R Tau) on the nervous system function and behavioral output, we pan-neuronally expressed *human mutant APP* (Aβ42) and *MAPT* (0N4R Tau), which resulted in a 20% and 40% reduction in climbing ability, respectively. Reduction in Ank2 expression likewise caused a 15% reduction in climbing, while Ank2 overexpression had no effect. All other gene combinations were found to significantly reduce climbing behavior.

### Overexpression of human mutant APP and MAPT genes and reduced Ank2 causes memory deficits

In order to determine the effect of the human mutant *APP* (Aβ42), *MAPT* (0N4R Tau) and *Drosophila Ank2* genes on memory we performed an olfactory shock assay [59-61] assessing memory at the 1 hour time point, which is considered to be intermediate memory. Olfactory shock memory is mediated by mushroom body (MB) neurons, we therefore drove expression of the genes using a promoter with broad expression in these neurons [59, 60]. We found MB expression of human mutant *APP* (Aβ42) or *MAPT* (0N4R Tau) caused a large reduction in memory (Fig. 5). *Ank2* is highly expressed in the adult MB as shown by *Ank2* reporter line (*R54H11-Gal4*) expression [62] and previous studies [57]. Reduction as opposed to overexpression of *Ank2* caused a similar reduction in memory. Co-expression of human mutant *APP* (Aβ42) and *MAPT* (0N4R Tau) or either AD gene with *Ank2-RNAi* caused a similar reduction in memory, suggesting that all three genes may act in the same pathway to regulate memory, an effect we localize to the MB. Overexpression of *Ank2* did not cause a significant reduction in memory, while co-expression of *Ank2* with human mutant *APP* (Aβ42) or *MAPT* (0N4R Tau) rescued memory to levels not significantly different to wild-type.

**Figure 5.**
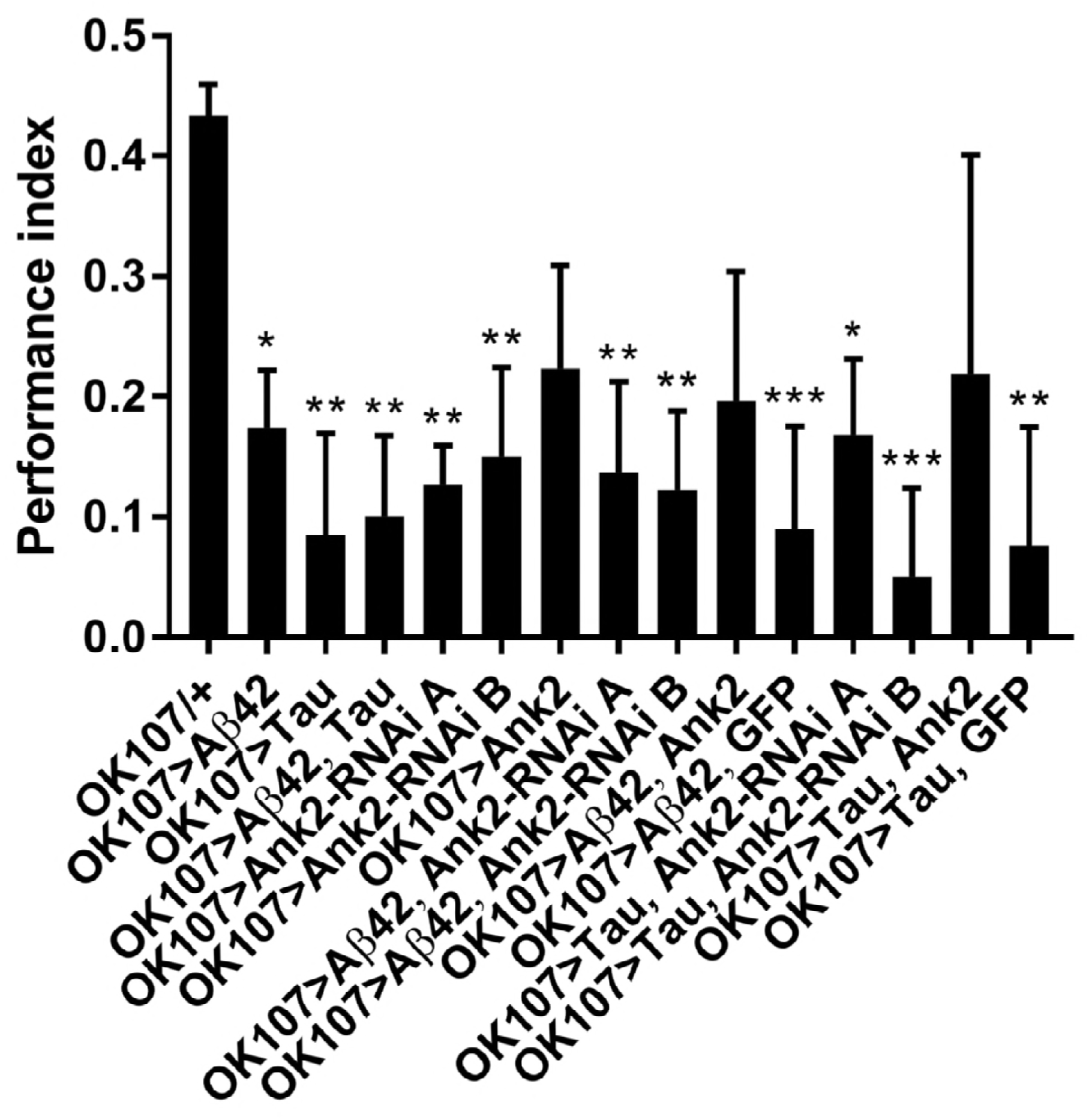
The effect of expression of human mutant *APP* (Aβ42), *MAPT* (Tau 0N4R) and *Drosophila Ank2* on memory. 1 hour memory was assessed using the olfactory shock-conditioning assay. The performance index of flies expressing AD transgenes throughout their mushroom body (MB) was compared to control (*OK107-Gal4/+*) were found to show a a significant reduction in memory using one-way ANOVA with Dunnett’s multiple comparisons, except genotypes overexpressing *Ank2*. Error bars are SEM. n≥4 groups of ∼100 flies.

In order for the flies to be able to perform the olfactory shock assay, the fly must be able to respond normally to shock and the odors used in the memory task. Therefore, we performed behavioral controls that showed that there was no significant difference between mutant genotypes and wild-type in terms of avoidance of shock (Fig. 6A), methylcyclohexanol (Fig. 6B) and octanol (Fig. 6C) odors [59-61]. Therefore, any mutant genotype showing a significant reduction in memory in Fig. 5 was a bona fide memory mutant.

**Figure 6.**
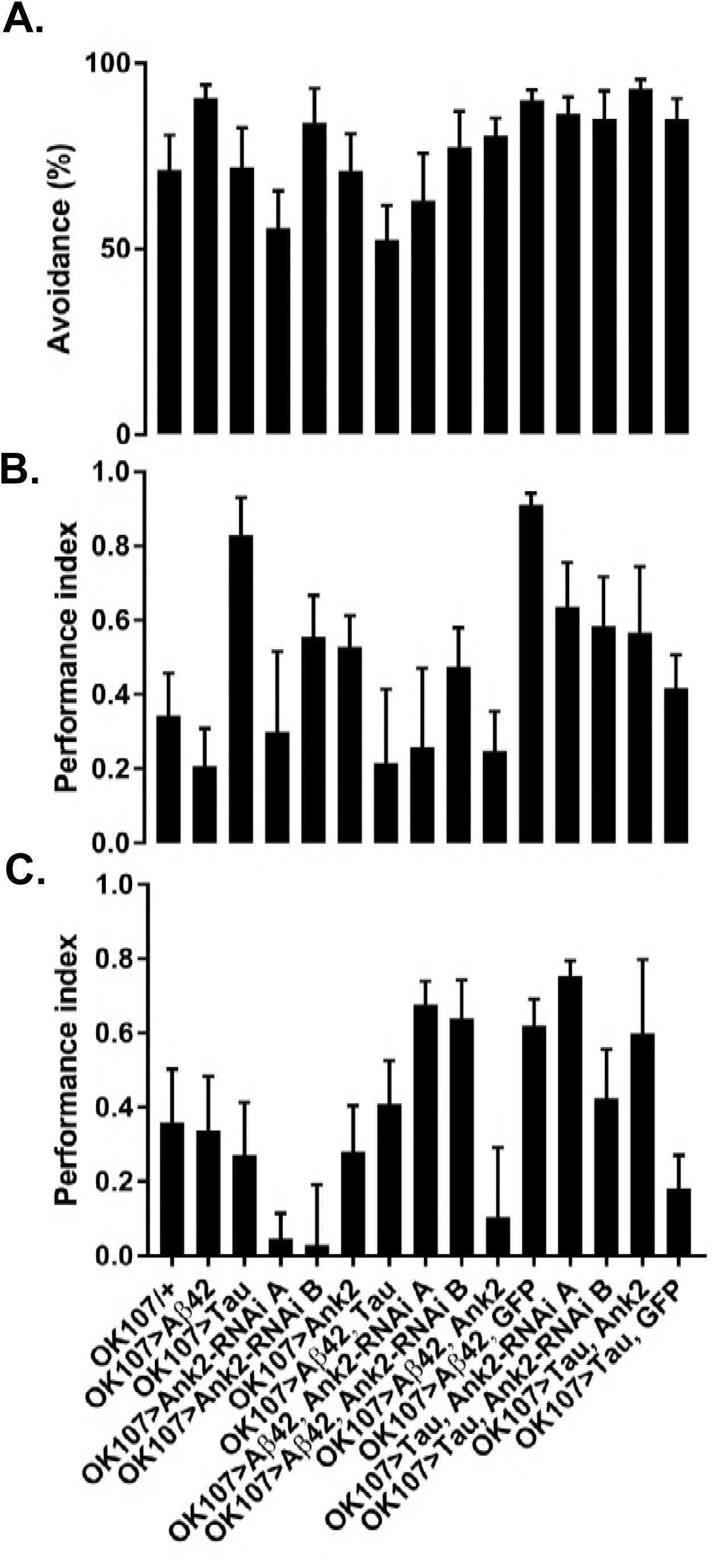
The effect of expression of human mutant *APP* (Aβ42), *MAPT* (Tau 0N4R) and *Drosophila Ank2* on response to negative reinforcement (shock) and olfaction. (**A**) The response of flies to negative reinforcement was quantified as % avoidance of shock, with experimental genotypes with AD genes expressed throughout their MB compared to control (*OK107/+*). The avoidance of these genotypes of flies to methylcyclohexanol (**B**) and octanol (**C**) was expressed as a performance index. All genotypes responded to odors and shock in a manner not significantly different from wild type using one-way ANOVA with Dunnett’s multiple comparisons. n≥4 groups of ∼50 flies.

### Mis-expression of human mutant APP (Aβ42), MAPT (0N4R Tau) and Ank2 decrease the neural excitability of memory neurons

Changes in neuronal excitability and Ca^2+^ signaling are thought to occur early in disease progression prior to neurodegeneration and are proposed to mediate early changes in behavior in AD, such as memory loss [24, 25]. Therefore, in order to determine how mis-expression of human mutant *APP* (Aβ42), *MAPT* (0N4R Tau) and *Ank2* may lead to changes in neuronal function and AD relevant phenotypes, we expressed the genetically encoded Ca^2+^ reporter, GCaMP6f in MB memory neurons and measured peak excitability in response to high [K^+^] solution that depolarizes neurons [60]. The axons and synaptic terminals of MB neurons form a pair of bilaterally arranged and symmetrically lobed structures which show low basal Ca^2+^ fluorescence levels (F0, Fig. 7A left panel) which in response to depolarizing high [K^+^] saline show a large peak in Ca^2+^ fluorescence levels (ΔF, Fig. 7A right panel). The increase in neuronal activity can be expressed as %ΔF/F0 time line plot (Fig. 7A bottom) showing high [K^+^] causes peak Ca^2+^ influx excitability that returns to baseline after washing. MB overexpression of human mutant *APP* (Aβ42), *MAPT* (0N4R Tau) or mis-expression of *Ank2* all caused a similar decrease in peak Ca^2+^ influx of memory neurons (Fig. 7B), suggesting a reduction in neuronal excitability.

**Figure 7:**
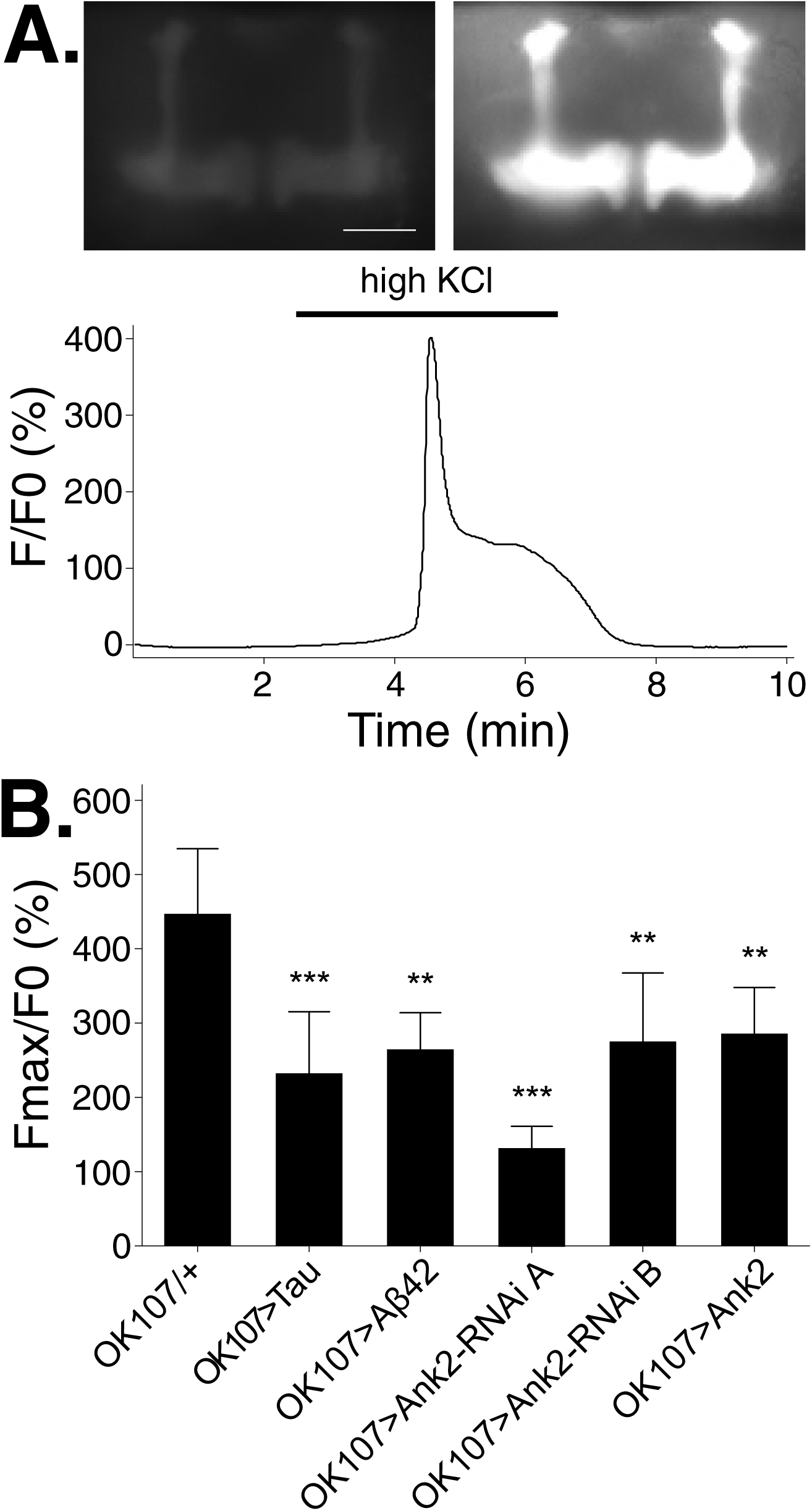
The effect of expression of human mutant *APP* (Aβ42), *MAPT* (Tau 0N4R) and *Drosophila Ank2* on activation of mushroom body neurons. **(A)** Exemplary control MB expressing the calcium reporter GCaMP6f shows a big increase in relative fluorescence in response to elevated KCl (100 mM, bath-applied as indicated by bar). Images show the same brain before and during KCl application; scale bar 50μm. **(B)** Quantitative analysis of the maximal response for the indicated genotypes shows a reduced responsiveness of all mutants. Data was analyzed with one-way ANOVA with Tukey’s *post hoc* and error bars are Standard Deviation. n≥5 flies.

## DISCUSSION

In this study we have generated and characterized the first animal model based on mis-expression of the AD EWAS nominated *ANK1* gene ortholog in *Drosophila.* Altered DNA methylation of human *ANK1* occurs in AD brains, especially in regions that show AD pathology, such as the entorhinal cortex [6, 7]. Here, we found reduction in expression of the fly ortholog of human *ANK1,* which is *Drosophila Ank2*, in areas of the brain responsible for memory (the MB), caused memory loss similar in magnitude to that caused by overexpression of human mutant *APP* resulting in an oligomerizing form of Aβ42 or *MAPT* resulting in the 0N4R isoform of Tau particularly associated with the disease. In flies, overexpression of the fly *Ank2* ortholog of the *ANK1* gene, did not result in degeneration of the eye, locomotor or memory defects but did lead to mild shortening of lifespan (by ∼10%) and reduced MB excitability. Interestingly *Ank2* overexpression in flies expressing human mutant *APP* (Aβ42) or *MAPT* (0N4R Tau), was found to return memory performance to a level statistically indistinguishable from wild-type. This suggests that *Ank2* may be acting in a similar pathway as Aβ42 or Tau to affect memory processing in the MB. Likewise, increasing *Ank2* expression reversed the degeneration of the eye caused by *MAPT* (0N4R Tau) overexpression in the eye. Therefore, altering the expression of *ANK1,* by targeting DNA methylation, may be considered a potential therapeutic strategy for treating AD.

In addition to memory loss, animals with reduced neuronal *Ank2* also recapitulated the shortening of lifespan seen in those with AD, again a phenotype seen in flies overexpressing human mutant *APP* (Aβ42) or *MAPT* (0N4R Tau). Co-expression of human mutant *APP* (Aβ42) and *MAPT* (0N4R Tau) caused a further reduction in lifespan, suggesting that the two molecules may act in partially non-overlapping and therefore additive pathways that lead to the pathology causing the flies to die early. A similar effect on lifespan has been reported previously for another form of *APP* that resulted in a non-oligomerizing version of Aβ42 and another form of *MAPT,* resulting in the 1N4R isoform of Tau, however the longevity assays were only run for roughly half the lifespan of the flies [40]. We also found a reduction in locomotion in flies expressing human mutant *APP* (Aβ42), *MAPT* (0N4R Tau) and reduced *Ank2*. Interestingly, a recent study has shown that *ANK1* hypermethylation is observed in PD, a movement disorder, that can also be associated with dementia [10]. In the eye expression of human mutant *APP* (Aβ42), *MAPT* (0N4R Tau) were both neurotoxic causing degeneration of the fly eye photoreceptor neurons. Co-expression caused a further reduction implying the two act in separate and additive pathways to cause neurotoxicity and neuronal death. Reduction in *Ank2* did not cause degeneration of the eye suggesting that *Ank2* may affect neuronal function independent of degeneration. This likely to be via changes in neuronal excitability, as we saw when the gene was misexpressed in MB neurons probably via ion channels, receptors or exchangers with which the adaptor molecule, ANK interacts with [11]. We found misexpression of human mutant *APP* (Aβ42), *MAPT* (0N4R Tau) and *Ank2* misexpression all reduced the peak Ca^2+^ response of MB neurons, a decrease in excitability likely to contribute to the memory deficits of these flies.

In *Drosophila, Ank2* has been shown to be important for synaptic plasticity and stability [54-56, 63, 64] and is involved in a glia mediated pathway that causes degeneration of motor neurons [54]. Therefore it is possible that reduction of *Ank2* may only cause degeneration in certain types of neurons in the eye. Furthermore, because human *ANK1* has also been shown to be misexpressed in glia in the AD brain [65], it is also possible that neurodegeneration results from misexpression of *Ank2* in glia.

In summary, we have made the first animal model of a gene implicated in AD that was nominated from EWAS. We found that mis-expression of the fly orthologue of this gene, *Ank2*, in neurons caused a range of AD relevant phenotypes such as shortened lifespan, memory loss and changes in neuron excitability similar to those resulting from human Aβ42 and 0N4R Tau. Furthermore increasing Ank2 expression reverses the memory loss and degeneration caused by overexpression of human mutant *APP* (Aβ42) and *MAPT* (0N4R Tau), suggesting that targetting *ANK1* may represent a new therapeutic target for the treatment of AD.

## MATERIALS AND METHODS

### *Drosophila* genetics

### *Drosophila* genetics

Flies were raised at a standard density with a 12 h:12 h light dark (LD) cycle with lights on at ZT 0 (Zeitgeber time) on standard *Drosophila* medium (0.7% agar, 1.0% soya flour, 8.0% polenta/maize, 1.8% yeast, 8.0% malt extract, 4.0% molasses, 0.8% propionic acid and 2.3% nipagen) at 25°C. The following flies used in this study were previously described or obtained from the Bloomington and Vienna fly stock centers: Wild type control was *Canton S w-* (*CSw-*) (gift from Dr Scott Waddell). Experimental genotypes were *elav-Gal4* (Bloomington stock center line number BL8760), *OK107-Gal4* (BL854), *GMR-Gal4* (BL9146), *uas-human MAPT (TAU 0N4R) wild-type* (gift from Dr Linda Partridge) [36, 66], *uas-human tandem Aβ42-22 amino acid linker-Aβ42* (gift from Dr Damian Crowther) [29], *uas-GFP* (gift from Dr Mark Wu), *uas-GCaMP6f* (BL42747 [67]), *uas-Ank2-RNAi* line A (BL29438), *uas-Ank2 RNAi* (Vienna *Drosophila RNAi* Center stock number VDRC107238 and KK10497) and *uas-Ank2* (gift from Dr RR Dubreuil) [53], *uas-Ank1-RNAi* line A (VDRC25946) and *uas-Ank1-RNAi* line B (VDRC25945).

### Survival assay

Approximately two days after eclosion ten mated females were transferred to a vial containing standard food and maintained at 25°C throughout. Deaths were scored every two days and then transferred to a fresh food vial [66]. Data was presented as Kaplan-Meier survival curves with statistical analysis performed using log-rank tests to compare survival between genotypes. All statistical tests were performed using Prism (GraphPad Software Inc., La Jolla, CA, USA).

### Eye degeneration assay

Adult flies of designated age were anesthetized by CO_2_ prior to immersion in ethanol in order to euthanize the fly to prevent any further movement during image capture [40]. The eyes were imaged with a Zeiss AxioCam MRm camera attached to a stereomicroscope (Zeiss SteREO Discovery.V8, up to 8× magnification). Surface area was quantified using Zeiss Zen software and normalized to the mean size of the wildtype control. One-way ANOVA with Dunnett’s multiple comparisons was used to analyze data.

### Climbing assay

Ten flies were collected and given 1hour to acclimatize to the test vial in an environmentally controlled room at 25°C and 70% humidity. Using the negative geotaxis reflex of *Drosophila*, flies were gently tapped to the bottom of a 7.5cm plastic vial and the number of flies that crossed a line drawn 2 cm from the top of the tube in 10 seconds was counted [27]. One-way ANOVA with Dunnett’s multiple comparisons was used to analyze data.

### Memory

1hr memory and sensory controls were performed as previously described using the olfactory-shock aversive conditioning assay [59-61]. Experiments were performed with groups of 30-50 flies of a given genotype, in a T-maze apparatus housed in an environmentally controlled room at 25°C and 70% humidity under dim red light. Flies were exposed to either 4-methylcyclohexanol (MCH, Sigma, ∼1:400) or 3-octanol (OCT, Sigma, ∼1:100) diluted in mineral oil (Sigma) paired with 1.5 second pulses of 60 volt electric shock interspersed with 3.5 second pauses from shock for a minute. After 30 seconds of fresh air the flies were exposed to the reciprocal order without shock for another minute. After a 1 hour rest, memory was assessed by transferring the flies to a choice point of the T maze, with one arm containing the shock paired odor and the other the non-shock paired odor, flies showed learning by avoiding the shock paired odor (i.e. correct flies).

Performance index (PI) = (number of correct flies – number of incorrect flies)/total number of flies.

To eliminate odor bias, the assay was performed with two groups of flies (30-50 flies in each), one shocked with MCH and then the other shocked with OCT. The average of taken of the two groups to give an n=1 PI value. Control experiments were performed to show that the different genotypes of flies could responded two MCH, OCT and shock alone. One-way ANOVA with Dunnett’s multiple comparisons was used to analyze data.

### Calcium imaging

Calcium imaging was performed using a genetically encoded GCaMP Ca^2+^ reporters and adapting previously published protocols [60, 67-70]. Adult flies of the indicated genotypes using *OK107-Gal4* and *uas-GCaMP6f* were collected between 2 and 5 days post eclosion, decapitated and the brain dissected in extracellular saline solution containing (in mM): 101 NaCl, 1 CaCl_2_, 4 MgCl_2_, 3 KCl, 5 glucose, 1.25 NaH_2_PO_4_, 20.7 NaHCO_3_, pH adjusted to 7.2. Brains were placed ventral side up in the recording chamber, secured with a custom-made anchor and continuously perfused with aerated saline. To activate neurons, high concentration KCl (100 μM in saline) was bath applied through the perfusion system for 4 min and then washed out. The calcium fluorescence signal was acquired using a CCD camera (Zeiss Axiocam) and a 470 nm LED light source (Colibri, Zeiss) on an upright Zeiss Examiner microscope with a 20x water immersion lens, recorded with ZEN (Zeiss, 4 frames/sec) and plotted with Microsoft Excel. One-way ANOVA with Dunnett’s multiple comparisons was used to analyze data.

## ACKNOWLEGMENTS

We thank Drs: Damian Crowther, Ronald Dubreuil, Linda Partridge, Scott Waddell and Mark Wu for *Drosophila* stocks and Dr Owen Peters for helpful comments on the manuscript. This work was supported by a GW4 accelerator (GW4-AF2-002) award to Drs: Katie Lunnon, Jonathan Mill, Jon Brown, Nick Allen, Vasanta Subramanian and James Hodge, as well as by Alzheimer’s Society undergraduate grants to Dr James Hodge. The authors declare no competing financial interests.

